# Hydrogen-deuterium exchange reveals a dynamic DNA binding map of Replication Protein A

**DOI:** 10.1101/2020.09.04.283879

**Authors:** Faiz Ahmad, Angela Patterson, Jaigeeth Deveryshetty, Jenna Mattice, Nilisha Pokhrel, Brian Bothner, Edwin Antony

## Abstract

Replication Protein A (RPA) binds to single-stranded DNA (ssDNA) and interacts with over three dozen enzymes and serves as a recruitment hub to coordinate most DNA metabolic processes including DNA replication, repair, and recombination. RPA binds ssDNA utilizing six oligosaccharide/oligonucleotide binding (OB) domains within a heterotrimeric complex of RPA70, RPA32 and RPA14 subunits. Based on their DNA binding affinities they are classified as high versus low-affinity DNA binding domains (DBDs). However, recent evidence suggests that the DNA-binding dynamics of DBDs better define their roles. Utilizing hydrogen-deuterium exchange mass spectrometry (HDX-MS) we assessed the contacts and dynamics of the individual domains of human RPA to determine the landscape of conformational changes upon binding to ssDNA. As expected, ssDNA interacts with the major DBDs (A, B, C, and D). However, DBD-A and DBD-B are dynamic and do not show robust DNA-dependent protection. DBD-C displays the most extensive changes in HDX, suggesting a major role in stabilizing RPA on ssDNA. DNA-dependent HDX kinetics are also captured for DBD-D and DBD-E. Slower allosteric changes transpire in the protein-protein interaction domains and the linker regions. We propose a dynamics-based DNA binding model for RPA utilizing a dynamic half and a less-dynamic half.

## INTRODUCTION

In eukaryotes, the single-strand DNA (ssDNA)-binding protein replication protein A (RPA) is essential for most DNA metabolic processes including DNA replication, repair, recombination, and telomere maintenance. RPA binds to ssDNA with high affinity (K_D_ >10^−10^M) (1) and protects it from degradation by exo- and endonucleases. Formation of RPA-ssDNA complexes triggers the ATM/ATR cellular DNA damage checkpoint response (2)(3). RPA physically interacts with over three dozen DNA processing enzymes and recruits them to the site of DNA damage. Finally, RPA hands-off the DNA to these enzymes and correctly positions them on complex DNA structures to facilitate their respective catalytic activity. Plasticity for such multiplexed functional roles for RPA can be ascribed to its flexible structure and its corresponding context-dependent interactions with DNA.

RPA is a heterotrimeric complex composed of RPA70, RPA32, and RPA14 subunits (also termed RPA1, RPA2, and RPA3, respectively). Structurally, it can be further partitioned into six oligonucleotide/oligosaccharide-binding folds (OB-folds; denoted A-F; Figure 1a). RPA70, the largest subunit of the heterotrimer, is composed of OB-folds F, A, B, and C. RPA32 consists of an N-terminal phosphorylation region, OB-fold D, and a C-terminal winged-helix domain (wh). Finally, RPA14 harbors OB-fold E (Figure 1a). The heterotrimer is constitutively held together by the trimerization core (Tri-C) which is mediated by interactions between OB-folds-C, D and E (Figure 1a) (4)(5). RPA mediates protein-protein interactions with over three dozen proteins primarily through two dedicated protein-interaction domains (PIDs). For clarity, here we partition the OB-folds as DNA binding domains (DBDs) and protein interaction domains (PIDs) based on their established biochemical functions. Domains A, B, C, D and E are DBDs whereas F and wh are PIDs. The F-domain in RPA70 (PID^70N^) and the ‘wh’ domain in RPA32 (PID^32C^) mediate direct physical interactions to most RPA interacting protein (RIPs). Some RIPs are composite binders and can interact with multiple sites on RPA. DBDs and PIDs are connected by flexible linkers of varying lengths (79 aa F-A linker; 10 aa A-B linker; 13 aa B-C linker; 32 aa D-wh linker) and thus allow RPA to adopt various conformations on ssDNA (Figure 1a).

**Figure 1.**
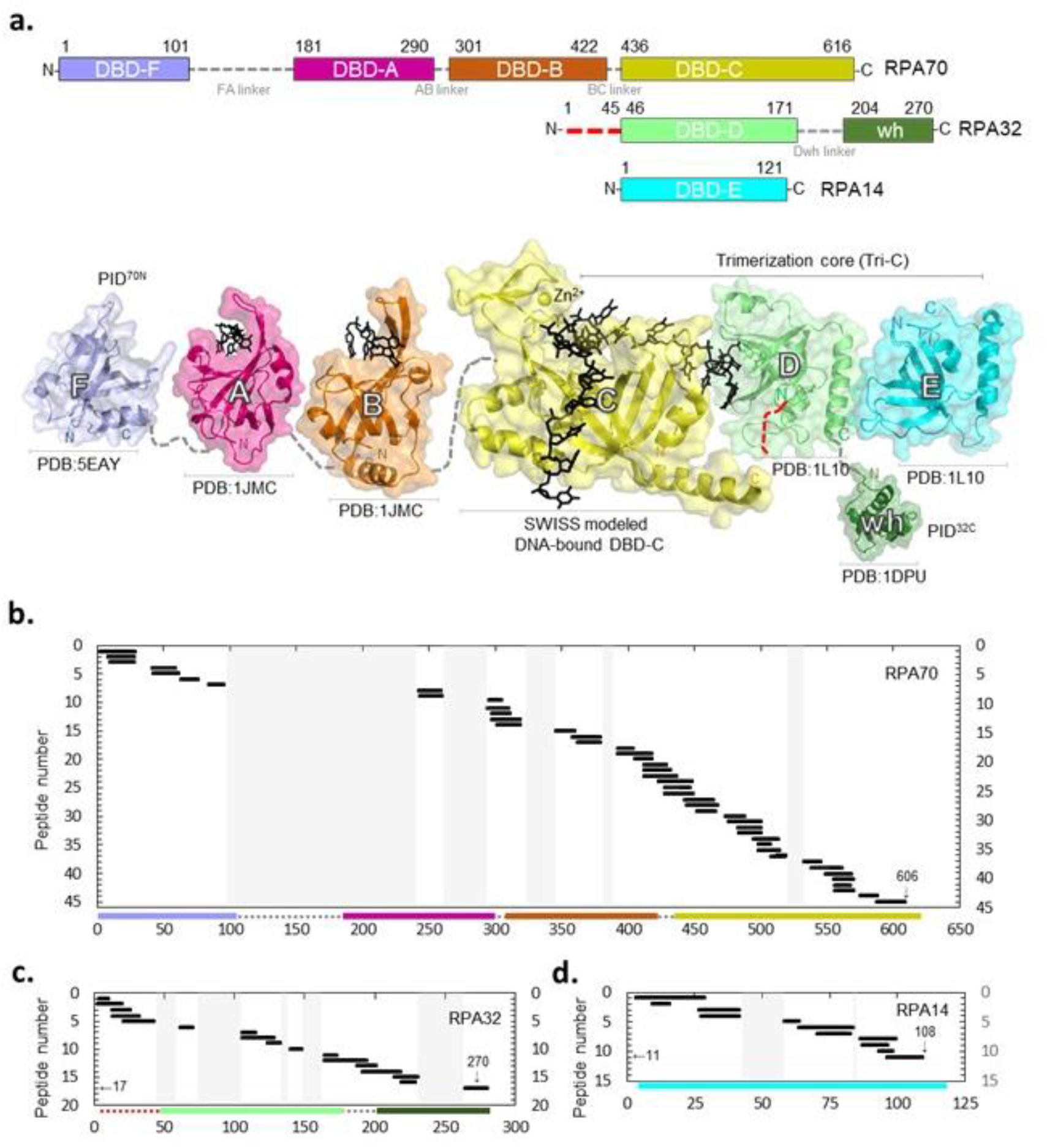
Sequence coverage of the DNA binding and protein interaction domains of RPA. **a**. Top - Domain composition of human RPA70, RPA32, and RPA14 subunits and their respective linkers. The red dotted line in the N-terminus of RPA32 denotes the region of extensive phosphorylation. Bottom – The crystal structures of the individual domains of RPA are depicted and presented to correspond to the panels on top. DNA is modeled onto DBD-C using SWISS-MODEL with the *Ustilago maydis* structure as reference. Peptides identified in the MS analysis corresponding to the three RPA subunits are shown. **b**. Shows 45 peptides from RPA70 providing 52% sequence coverage. The 17 peptides in **c**. corresponds to RPA32 providing 50% sequence coverage. **d**. Showcases 11 peptides from RPA14 with 49% sequence coverage. The y-axis indicates the peptide fragment number and the x-axis denotes peptides arranged in order of increasing residue numbers from the N-terminus of each subunit as denoted in Supplemental Table 1.

Although RPA binds to ssDNA with high affinity, it must be removed by RIPs that bind ssDNA with much lower affinity. This is achieved by the differential DNA binding properties of individual DBDs. Several structural and biochemical studies have identified the DNA binding regions and properties of RPA (5)(6). Human RPA displays multiple DNA binding modes depending on the NaCl concentration and denoted by their occluded site-size. This varies from 22 nucleotides (nts) in conditions below 50 mM NaCl to 28-30 nts at higher salt concentrations of up to 1.5 M NaCl (7). A sequential DBD binding model has been proposed and posits that smaller occluded site-sizes are likely due to the higher affinity ssDNA binding properties of DBDs-A and B, whereas binding of all DBDs drive the 30 nt mode. Support for this model arises from DNA binding studies of the isolated individual DBDs, where DBDs-A and B were defined as the high affinity binders with DBDs-C and D as low affinity ssDNA binding domains (8)(9)(10). Moreover, both DBD-E and PID^70N^ possess very weak DNA binding properties (4)(11)(12)(13)(14)(15). The transition between the DNA binding modes potentially drives the diffusion of RPA on ssDNA and transiently provides access to buried DNA for RIPs to latch on (7). The DNA-bound crystal structure of *Ustilago maydis* RPA provides an excellent snapshot of how the DBDs of RPA are arranged on ssDNA. DBDs-A, B, C and D engage the DNA and parlay 5′-3′ directionality with DBD-A residing towards the 5′ end (4)(16)(17)(18)(19)(20)(21). PID^70N^ and PID^32C^ are not present in the structure, and DBD-E does not contact the DNA. Most of the intervening linker regions are also disordered (*i*.*e*. flexible; electron density not visualized) in the structure.

Due to the inherent conformational flexibility, obtaining an apo-structure of full-length RPA is challenging. This has resulted in limited capture of RPA structural transitions (apo to DNA bound state) using methods as XRC (X-ray crystallography), SAXS (small angle X-ray scattering) or cryoEM (cryo electron microscopy). XRC or NMR spectroscopy high resolution structures have been solved for all DBDs of RPA (22)(23). SAXS studies have shown PID^70N^ to be flexibly linked to DBDs-A and B and has no impact on the ssDNA binding properties of these DBDs (24). Recent cryoEM studies of *Saccharomyces cerevisiae* RPA on longer ssDNA (100 nts) showcases the interplay between the DBDs when multiple RPA molecules are bound (23). In this structure, DBDs-A and -B are disordered compared to Tri-C. This finding contradicts the high-affinity binding label ascribed to DBDs-A and B. Furthermore, the structure shows that DBD-A from one RPA molecule physically interacts with DBD-E of the neighboring RPA molecule. Such an arrangement opens up segments of ssDNA bound between two RPA molecules, likely serving as DNA binding regions for RIPs. Finally, PID^70N^ and PID^32C^ are disordered in the cryoEM structure further solidifying the fact that they are primary contributors for interacting with RIPs and do not contribute much to the DNA binding properties of RPA, although potential allosteric effects cannot be ruled out.

Recent single-molecule experiments support an alternative DNA binding dynamics-based model for RPA. Each DBD, or a subset of DBDs have defined on-off properties on ssDNA that dictate formation of specific RPA conformations on DNA (4)(20)(21). DBD-A and DBD-D have four binding states on DNA with lifetime of each state differing between the two DBDs. Bulk fluorescence kinetic experiments have shown conditional displacement of DBD-A (and likely DBD-B) by Tri-C (23)(25)(26). The dynamics and arrangement of individual DBDs can also be altered by post-translational modifications and interactions with RIPs. Phosphorylation at S178D, a position on the FA-linker adjacent to DBD-A ablates DBD-A binding to ssDNA and alters its positioning with respect to the neighboring DBDs (23). Similarly, the presence of Rad52, a mediator protein that functions in homologous recombination, alters the number of states formed by DBD-D on ssDNA (25). Thus, knowledge of the kinetic stability of specific DBDs on ssDNA might help in establishing whether each of the DBDs is either stable or dynamic. We use the term *dynamic* or *dynamics* to imply residence time on DNA. Since such experiments have to be performed using the full length RPA protein, we here utilized time-resolved hydrogen-deuterium exchange mass spectrometry (HDX-MS) to capture the conformational changes within human RPA as a function of time upon binding to ssDNA.

HDX-MS is ideal for studying protein folding and its conformational dynamics (27)(28)(29)(30). HDX-MS measures the rate of amide-H exchange to report on changes in the local environment. D_2_O exchange rates are shaped by dynamics, H-bonding, secondary structure, and solvent exposure. Increased rates of D_2_O exchange result from greater solvent accessibility, disruption in backbone H-bonding, or increased dynamics. Decreased D_2_O exchange rates suggest less solvent accessibility or local stabilization (*i*.*e*. H-bonding and/or formation of secondary structure). As such, exchange rate disruptions are valuable indicators of protein allostery, dynamics and interfaces (31). In recent years, HDX-MS has gained attention as powerful tool to study protein-DNA binding interactions (27)(30)(32)(33). Here, using HDX-MS we identify the contacts between the various DBDs, linkers, and PIDs of human RPA upon complex formation with ssDNA. Based on the data, we propose an alternate dynamics-based model for RPA: DBD-A, DBD-B, and PID^70N^ (all in RPA70) serve as the *dynamic half* of RPA. Tri-C, consisting of DBD-C (RPA70), DBD-D (RPA32) and DBD-E/RPA14, function as the *less dynamic half* with DBD-C serving as the anchor that likely controls the residence time of RPA on ssDNA.

## MATERIAL AND METHODS

### HDX-MS

Human RPA was purified as described (25). RPA and DNA (dT)_30_ were mixed as 1:1 molar equivalents (3.9 mg/mL RPA) and diluted 1:10 into deuterated reaction buffer (30 mM HEPES, 200 mM KCl pD 7.8 in D_2_O). Undeuterated controls (zero time points) were diluted 1:10 into non-deuterated buffer. 10 µL of the reaction was removed at each time point (0.008, 0.05, 0.5, 3, and 24h) and quenched on ice by adding 60 µL of 0.75% formic acid (FA, Sigma) pH 2.5 containing 0.25 mg/mL porcine pepsin (Sigma). Each sample was digested for two minutes with vortexing every 30 seconds and then flash frozen in liquid nitrogen. Samples were stored in liquid nitrogen until liquid chromatography mass spectrometry (LC-MS) analysis. All reactions were performed in triplicate.

LC-MS analysis of RPA was performed on a 1290 UPLC series chromatography stack (Agilent Technologies) coupled directly to a 6538 UHD Accurate-Mass QTOF LC/MS mass spectrometer (Agilent Technologies). Before electrospray−time-of-flight (ESI-TOF) analysis, peptides were separated on a reverse phase (RP) column (Phenomenex Onyx Monolithic C18 column, 100 × 2 mm) at 1 °C using a flow rate of 500 μL/min under the following conditions: 1.0 min, 5% B; 1.0−9.0 min, 5−45% B; 9.0−11.8 min, 45-95% B; 11.80−12.0 min, 5% B; solvent A = 0.1% FA in water (Thermo-Fisher) and solvent B = 0.1% FA in acetonitrile (ACN, Thermo-Fisher). Data were acquired at 2 Hz over the scan range 50−1700 m/z in positive mode. Electrospray settings were as follows: nebulizer set to 3.7 bar, drying gas at 8.0 L/min, drying temperature at 350 °C, and capillary voltage at 3.5 kV.

Peptides were identified as outlined in *Berry et al*. (28) using MassHunter Qualitative Analysis version 6.0 (Agilent Technologies), Peptide Analysis Worksheet (PAWs, ProteoMetrics LLC), and Peptide Shaker version 1.16.42 paired with Search GUI version 3.3.16 (Compomics). Deuterium uptake was determined using HDExaminer version 2.5.1 (Sierra Analytics).

### Data analysis

Average relative deuterium uptake difference (ΔHDX) averaged over all HDX incubation period was calculated using equation (34).

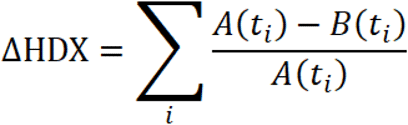

The ΔHDX between bound and free RPA was calculated by the following equation, where A is the deuterium uptake for sample A (RPA+DNA) at one time point (ti), and B is the deuterium uptake for sample B (free RPA) at the same time point (ti). Summation of ΔHDX at all five time point together gives ΔHDX for a peptide (Figure 2d-2f). Mean of the deuterium uptake value of a peptide was used to generate a heatmap and plotted using GraphPad Prism 8.4.1 software. To draw crystal structure PyMOL 4.6 was used. The uptake curve for each peptide was plotted using a non-liner/non-logarithmic scale using MS-Excel. Each data point represents the mean value, and error bars represent the ±SD of mean for biological replicates. Peptides with a SD> 0.5 were excluded from this study. To compare the difference at time points with ±DNA statistical significance were determined. *, the first incubation time point that shows statistical difference are indicated in the linear plot. No significance is indicated as ‘ns’ and p<0.05 was considered statistically significant (* p-value < 0.05, ** p<0.01, ***p<0.001, two-tailed, unpaired t test). In the full-length RPA plasmid, RPA32 is his-tagged, thus HDX peptide 17 had an extra 6xHis at its C-terminus i.e., TDAEHHHHHH.

**Figure 2.**
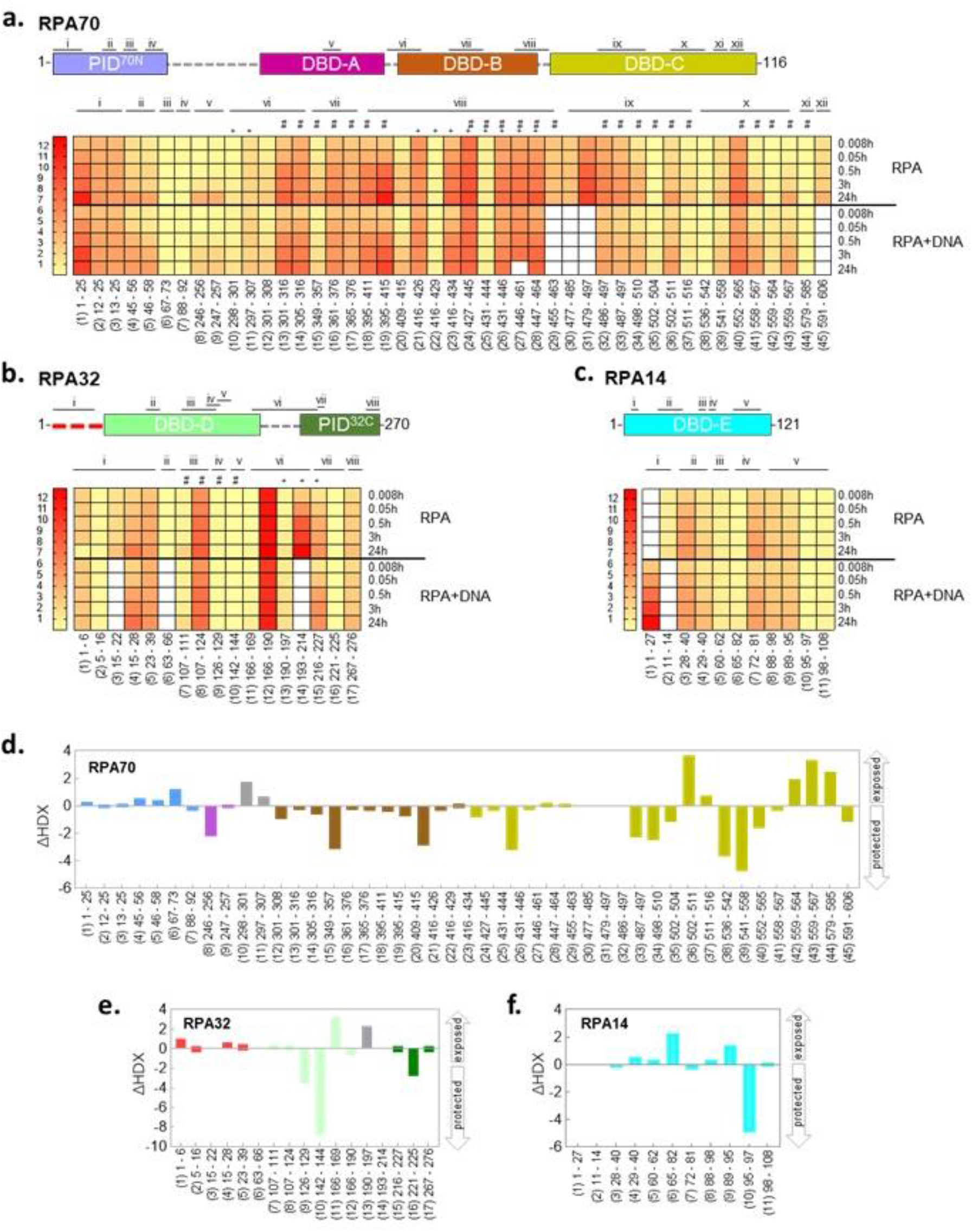
Global deuterium uptake profiles and DNA induced changes in RPA. Heat map representing the deuterium content of **a**. RPA70, **b**. RPA32, and **c**. RPA14 peptides in unbound and DNA bound states as a function of time. Each row represents a timepoint and each column represents a peptide. The heat map gradients are color coded yellow (low deuterium uptake) to red (high deuterium uptake). The white boxes depict absence of a corresponding dataset for specific peptides under one condition (presence or absence of DNA). # Denotes peptides in DNA binding sites, + marks peptides from the linker regions. To map the peptide location in RPA70, RPA32 and RPA14, peptides in same region were grouped from i to xii. Binding to ssDNA leads to varying patterns of protection by HDX-MS in **d**. RPA70, **e**. RPA32, and **f**. RPA14. The arrows denote whether the peptides were protected or exposed to solvent upon DNA binding. The data show the cumulative difference in deuterium uptake (ΔHDX) in RPA over all five HDX timepoints.

## RESULTS

### HDX-MS generates a conformational landscape for DNA-induced changes in human RPA

To determine the ssDNA binding associated conformational changes in RPA, we performed HDX-MS experiments with human RPA in the absence and presence of ssDNA. A (dT)_30_ oligonucleotide was used to provide adequate binding sites for all the domains of RPA as the binding site-size for RPA is ∼22-28 nt (35). Since RPA binds to ssDNA stoichiometrically, we used an equimolar concentration of RPA and (dT)_30_. Given the size of RPA (1007 amino acids in total *i*.*e*. RPA70 - 616 aa, RPA32 - 270 aa, and RPA14 - 121 aa), we observe an excellent sequence coverage map of full-length RPA (Figure 1b-1d; and Supplemental Table 1). We identified 45 (RPA70), 17 (RPA32), and 11 (RPA14) different peptides covering RPA sequence (Supplemental Table 1). This provided an overall 51% sequence coverage (RPA70 – 318 of 616 aa ∼52%, RPA32 – 139 of 276 aa ∼50%, RPA14 – 59 of 121 aa ∼49%) with majority of the unreported sequence in the FA-linker and DBD-A domains (Figure 1b). For much of the complex, multiple overlapping peptides were identified. These were grouped and assigned Roman numerals for clarity (Figure 2). There are 12 groups for RPA70 (i – xii), 8 groups for RPA32 (i – viii), and 5 groups for RPA 14 (i – v) (Figures 2a, 2b and 2c), respectively.

RPA binds rapidly to ssDNA (*k*_on_ =1.1±0.6×10^8^ M^-1^s^-1^) (25). Therefore, we reasoned direct DNA-binding induced changes would occur rapidly, while diffusion and DBD rearrangement associated deuterium exchange would occur on a longer timescale. To get a better idea of such changes we performed the HDX-MS analysis as a function of time to obtain a direct readout of the kinetics of deuterium exchange in the absence and presence of ssDNA. We present the data in terms of the kinetics and patterns of HDX. In terms of the HDX kinetics, two prominent deuterium exchange patterns are observed which we classify as *fast* and *slow exchange*. Fast exchanges are deuteration that has occurred before the first time point captured in our experiments. RPA binding to ssDNA is diffusion limited; thus, the initial DNA-induced rapid conformational changes occur faster than our ability to capture them. Differences in exchange kinetics that occur only on a fast timescale, suggest rapid and stable binding to ssDNA. This is indicative of a lasting H-bond networks and a stable structure. Slow exchange is used here to describe deuterium exchange occurring at later time points. Slow exchange is a result of dynamic H-bond networks leading to solvent exposure. This infers slower on/off rates due to conformational changes and dynamics. Difference in HDX rates that occur over a 3 to 30 min interval are likely due to conformational rearrangements of the domains and other allosteric effects, which we have classified as *slow exchange*.

With respect to HDX patterns, we either observe a *protection* or *exposure* of certain regions of RPA upon binding to ssDNA. In our experiments, data is collected after mixing RPA with ssDNA. Since the binding is diffusion limited, the deuterium uptake or loss at the first time point (30 sec) likely reflects a direct DNA-induced change. If a domain forms a long-lived stable complex on DNA, a corresponding long-lived difference in deuterium exchange will be observed compared with RPA alone. We classify such a signal as *protection*. This can be thought of as protein surfaces that become buried upon DNA binding and do not come apart. If exchange continues at the minutes to hours timepoints, then those peptide regions are transiently exposed to solvent and are therefore ‘less’ protected or *exposed*, hence more dynamic.

The deuterium exchange data for all the peptides are summarized as a heat map where red and yellow indicate high and low deuterium uptake, respectively (Figures 2a-c). White boxes denote absence of data as one of the corresponding peptides were not identified in the MS analysis. Peptides 1 to 5, and 10 to 45 show the highest degree of deuteration and correspond to PID^70N^, DBDs -A, -B, and -C in RPA70, respectively (Figure 2a). For RPA32, peptides 7 to 8 and 11 to 14 in DBD-D show larger changes in deuteration when bound to DNA. Peptides 15 and 17, which are in the winged-helix domain (PID^32C^) show less exchange (Figure 2b). Thus, both protein interaction domains in RPA (PID^70N^ and PID^32C^) show minimal changes that transpire on a slower timescale, likely reflecting conformational changes driven indirectly after DNA binding to the DBDs. Peptides 5, 6 and 8 to 11 in DBD-E show moderate deuterium exchange, but the fast exchange kinetics coincide with the rate of DNA binding (Figure 2c). This is consistent with a direct interaction between DBD-E and DNA as suggested by crosslinking experiments (6)(36).

To construct a global landscape of the DNA induced deuterium exchange in RPA, we quantified the average relative deuterium uptake difference (ΔHDX) between the deuteration in the absence and presence of DNA (Figures 2d-f). The difference in deuteration for each DNA binding experiment relative to apo RPA is averaged over time for every peptide (Equation 1). The scale of the bars denotes the net change in deuterium uptake. A change in the upward direction denotes increased deuterium uptake (‘less’ protected or *exposed*) in the presence of DNA. Conversely, data movement in the downward direction denotes less uptake (*protected*) upon DNA binding. The ΔHDX data show robust deuterium exchange in DBD-C with additional changes in DBD-A and DBD-B as expected from known DBD-ssDNA interactions (Figure 2d). Similar ΔHDX data are observed in DBD-D (RPA32; Figure 2e) and in DBD-E (Figure 2f). DBD-E binding to DNA has been observed in biochemical studies (11), but not in the X-ray or cryoEM structures of RPA-DNA complexes. Our data show DNA-driven fast exchanges in DBD-E. ΔHDX describes the difference in deuterium incorporation under two distinctive conditions. Detailed analysis of the deuteration changes across the various domains of RPA are discussed below.

To best represent the data, we mapped the peptides identified in the MS analysis onto the available crystal structures of human RPA, and the regions corresponding to the peptides are highlighted in red (Figures 3, 4 and 5 and Supplemental Figures 1-7). Since a structure of full-length human RPA is not available, we constructed one for reference using the information available for the two halves of human RPA. We used the ssDNA-bound structure of DBD-A, and -B (PDB ID:1JMC) (37) and the structure of RPA Tri-C (PDB ID: 1L10) (4). However, the Tri-C structure is not in complex with DNA. Thus, we used the DNA-bound structure of *Ustilago maydis* RPA as reference and modeled DNA onto human RPA Tri-C using SWISS-MODEL (38). Our data below are presented from the perspective of two dynamic halves in RPA: PID^70N^, DBD A and DBD-B constituting the ‘*FAB half’*, and the ‘*Tri-C half*’ made of DBD-C, DBD-D, PID^32C^ and DBD-E.

**Figure 3.**
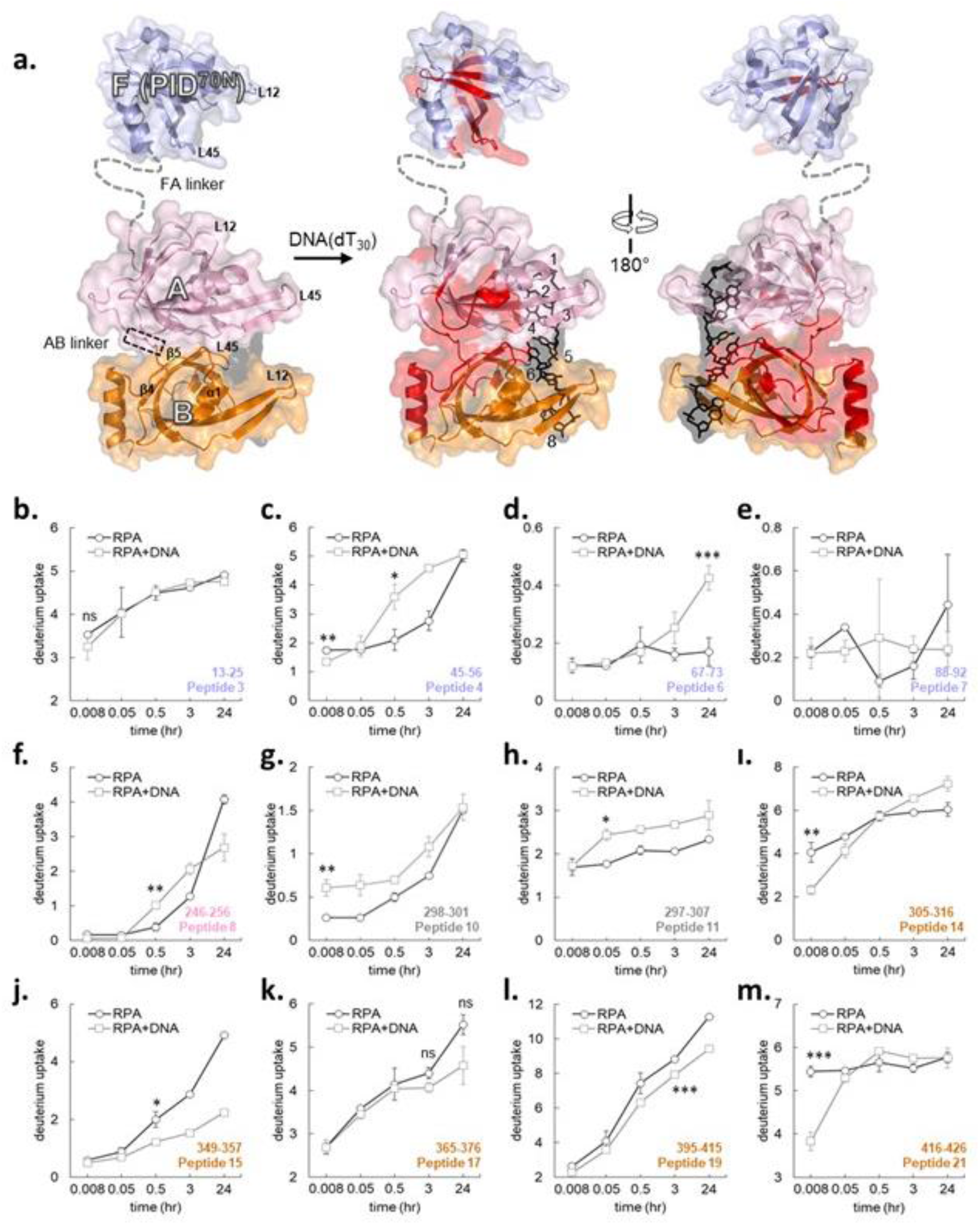
Time-resolved HDX-MS analysis of DNA-induced conformational changes in the FAB half of RPA. **a**. Peptides identified in the MS analysis are mapped onto the crystal structure of PID^70N^, DBD-A, and DBD-B. The F-AB and A-B linkers are denoted by dotted lines. **b-m**. Deuterium exchange profiles of individual peptides from the FAB half of RPA. Error bars reflect standard deviations, calculated as described in Material and Methods. *, the first incubation time point that shows statistical difference between free RPA and DNA bound RPA. No significance indicated as ‘ns’.

**Figure 4.**
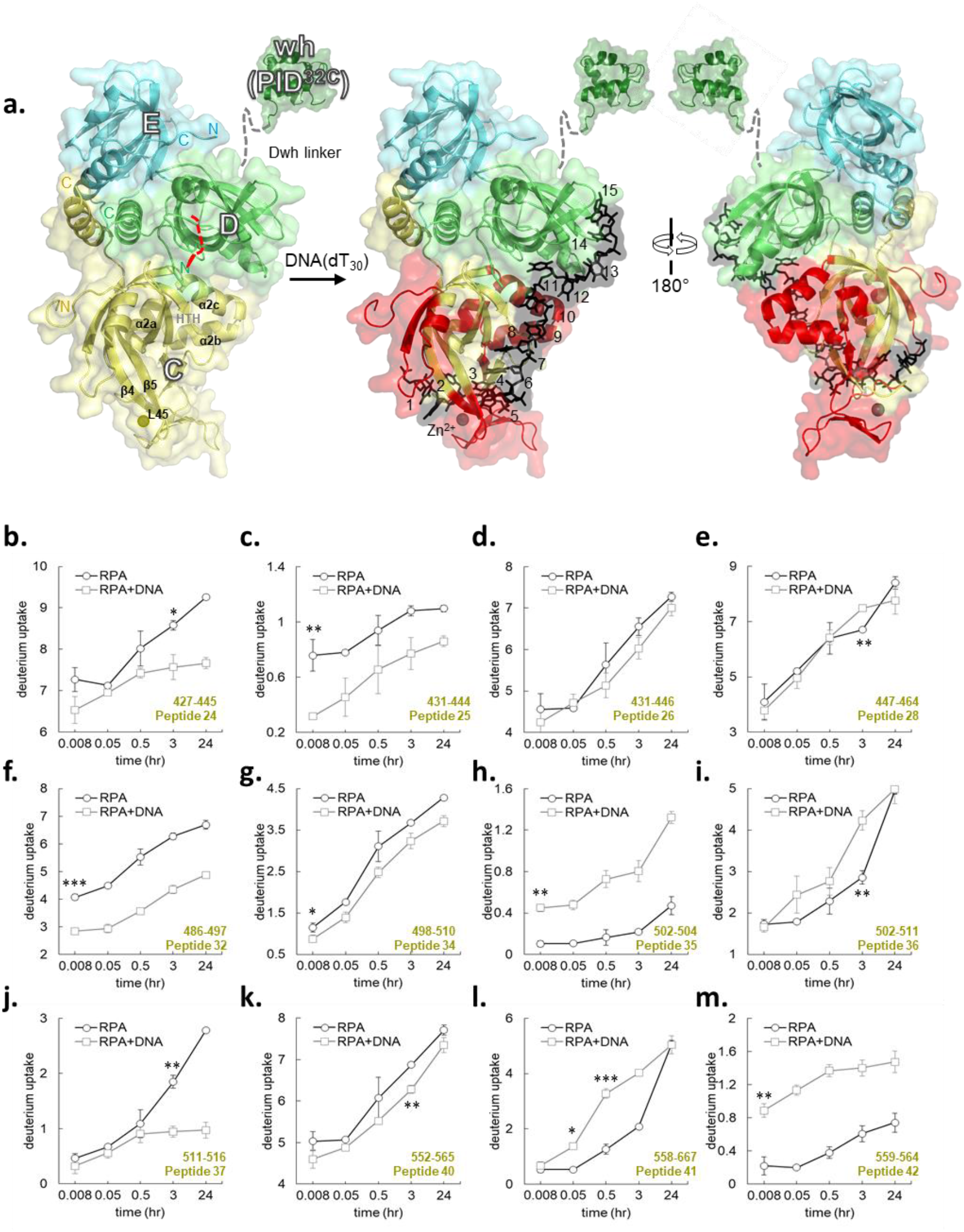
Time-resolved HDX-MS analysis of DNA-induced conformational changes in DBD-C. **a**. Crystal structure of the Tri-C core of human RPA composed of DBD-C, RPA32 and RPA14. The peptides identified the MS analysis corresponding to DBD-C are denoted in red. **b-m**. Deuterium exchange profiles of individual peptides from DBD-D are shown. Error bars reflect standard deviations, calculated as described in Material and Methods. *, the first incubation time point that shows statistical difference between free RPA and DNA bound RPA. No significance indicated as ‘ns’.

**Figure 5.**
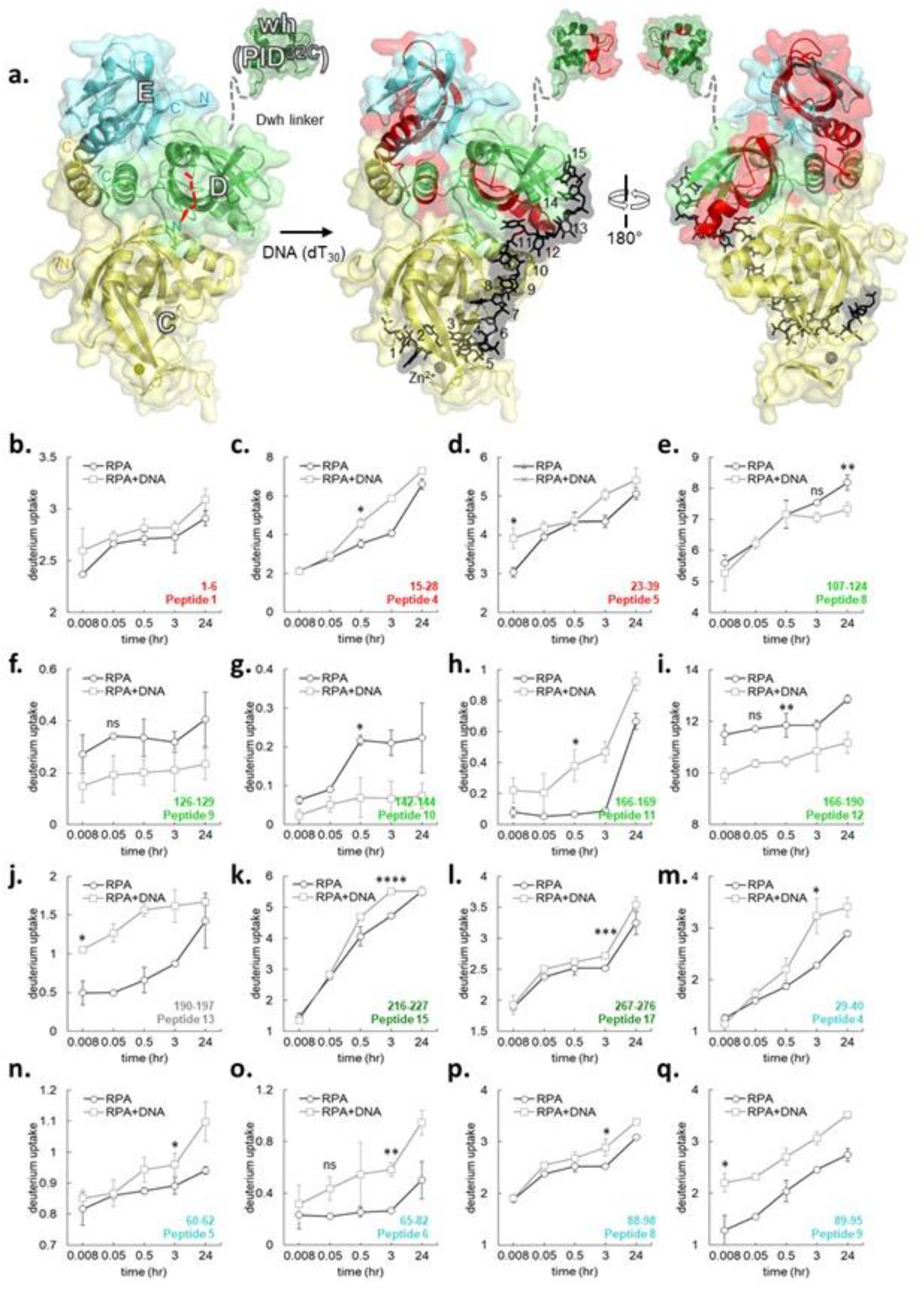
Time-resolved HDX-MS analysis of DNA-induced conformational changes in RPA32 and RPA14. **a**. Crystal structure of the Tri-C core of human RPA composed of DBD-C, RPA32 and RPA14. The peptides identified the MS analysis corresponding to RPA32 and RPA14 are denoted in red. **b-q**. Deuterium exchange profiles of individual peptides from RPA32 and RPA14 are shown. Error bars reflect standard deviations, calculated as described in Material and Methods. *, the first incubation time point that shows statistical difference between free RPA and DNA bound RPA. No significance indicated as ‘ns’.

### The FAB half of RPA shows highly dynamic ssDNA-dependent changes in HDX

PID^70N^, DBD-A and DBD-B reside in RPA70 and are connected by flexible linkers. The 79 aa FA-linker connects DBD-A to the PID^70N^ and the shorter 10 aa long AB-linker links DBD-A and -B (Figure 3a). DBD-A and -B are well-established DNA binding domains (K_D_ = 2 μM DBD-A; 20 μM DBD-B; 50 μM DBD-A+B) (4)(9)(10). Canonically, these DBDs are considered as the high affinity DNA binding domains of RPA. For PID^70N^ we observe changes on longer time scales accompanied by a slow decrease in protection upon DNA binding (Figures 3b-e and Supplemental Figures S1a-c). Peptides 4 and 5 in region ii are part of the basic cleft of the proposed weak DNA binding site of PID^70N^. Both peptides show similar total exchange with or without DNA, but the uptake rate is slower without DNA (Figure 3c and Figure S1c). Peptide 6 is protected without DNA, becoming more dynamic (exchange over a long timescale), with slow conformational changes when DNA bound (Figure 3d). These observations suggest that the conformational changes in PID^70N^ in the presence of DNA occur on a timescale well past the initial formation of the RPA-DNA complex.

For DBD-A, two overlapping peptides were observed, both showing limited HDX change. Peptides 8 and 9 are not in the DNA binding cleft, rather they are in the α2/β4 loop region located opposite to the cleft (Figure 3f, and Supplemental Figure 2b). Both peptides show different total exchange with or without DNA, but the uptake rate is slow without DNA. In the absence of DNA, peptides 10 and 11 from the AB-linker show slow and fast deuterium exchange overtime, respectively. Upon DNA binding an increase deuterium uptake is observed for both the peptides, however, the pattern was different (Figures 3g-h). The rate of exchange for peptide 10 was impacted at the first time point, whereas peptide 11 was impacted at later time points. This suggests that protection and dynamics are altered by DNA binding. Presumably, since the linker is located closer to the DNA binding cleft of DBD-B, greater deuterium uptake in these two peptides is likely directly tied to DNA binding (Supplemental Figures 2b-c). Overlay of apo and DNA bound DBD-A show overall conformational changes upon DNA binding (Supplemental Figure 2d).

DBD-B yielded better MS coverage with a total of 14 peptides (Figure 1b and Supplemental Table 1) and provides a clear picture of conformational changes. Surprisingly, the HDX changes in this domain were minimal upon DNA binding. Most peptides in this region (peptides 12, 13, 16, 18) do not show significant changes in HDX (Supplemental Figures 3a-d). Peptides 14 and 15 showed slower conformational changes upon DNA binding (Figure 3i-3j). Peptide 15 in DBD-B (β3-strand) shows slow deuterium uptake kinetics with significant (*p=0.016) decreased uptake (*protection*) at later time points with DNA. Regions within the DNA binding cleft (peptides 16, 17, 18, 19) have similar D_2_O exchange pattern, with a minor decrease initially that is more pronounced at later time points upon DNA binding (Figures 3k-3l and Supplemental Figures 3c-d). In the C-terminus of DBD-B (α2-helix; peptide 20), DNA binding leads to protection from D_2_O exchange (Supplemental Figure 3e). Peptide 21 shows significant (***p=0.0007) DNA-induced protection at the first time point (Figure 3m). Peptides corresponding to the BC-linker (peptides 21, 22 and 23) displays dynamic changes with DNA dependent protection (Supplemental Figures 3f-g). Along with the DNA induced conformational changes, HDX kinetics in several regions of DBD-B fluctuate between time points, especially within the DNA binding cleft. Comparison of apo (PDB:1FGU) and DNA-bound (PDB:1JMC) DBD-B structures reveals several regions undergoing conformational changes. Specifically, largescale changes occur in the L12 and L45 loop regions (Supplemental Figure 3h).

Our data support a dynamic model for DBD-A and DBD-B engagement in agreement with recent single molecule fluorescence studies (23)(25). DBD-A and DBD-B can adopt an array of conformations due to flexible and rotational effects at the flexible linker (21). Our HDX data is consistent with conformational flexibility of these domains. Residues 314 to 316 (peptide 13-14) and 395 to 401 (peptide 18-19) are at the interface between DBD-A and -B in the DNA bound structure and are solvent occluded. These residues show protection upon DNA binding (Figures 2d, 3i-3j, 3l, and Supplemental Figures 3b & d). Unfortunately, we do not have coverage data for DBD-A at the same interface. Nevertheless, our observations agree with the existence of various conformational states of DBD-A and DBD-B. There are limited contacts between DBD-A and -B with a flexible linker in apo structures which are brought together upon DNA binding (39). Time resolved deuterium exchange in the FAB half of RPA shows gradual increase in deuterium uptake that are not linear. Furthermore, the amplitude of deuterium uptake does not reach saturation (Figure 3, and Supplemental Figures 1-3). Such deuterium exchange profiles are characteristic of protein-DNA interactions that are dynamic with rapid on and off rates (32)(40). The dynamic DNA binding properties of the FAB half are in agreement with recent single molecule studies that showcase multi-step dynamic interactions between DBD-A, DBD-B and ssDNA (23)(25).

### Tri-C shows less dynamic ssDNA-dependent changes in HDX

The trimerization core (Tri-C) is composed of DBD-C (RPA70), DBD-D (RPA32), PID^32C^ and DBD-E (Figure 4a). Structural and biochemical evidence suggests that Tri-C works as a single constitutive unit and binds to 24 nts (4). DBD-C is connected to DBD-B by the BC-linker (13 amino acids) and can be envisioned as a hinge holding the two halves of the RPA together (Figure 1a). HDX analysis of DBD-C, the largest DNA binding domain in RPA, yields excellent coverage with total 24 peptides (Figure 1b and Supplemental Table 1). The HDX changes in DBD-C align well with expectations for a stable DNA binding domain. Most differences in deuterium uptake upon DNA binding in DBD-C occurred on fast timescales and were of significant magnitude (Figures 2a, 4 and Supplemental Figure 4). Part of the BC-linker and the N-terminus of DBD-C (peptides 24, 25, 26, and 28) consistently show deuteration differences at every time point with a noticeable decrease in deuterium uptake upon DNA binding (Figures 4b-e). The N-terminus is less protected on DNA binding (peptide 27, Figure S4a). The BC-linker in the *U. maydis* RPA structure assumes two β-turn structures (41). The first β-turn inserts into the DNA binding groove of DBD-B and is proposed to preclude DNA from adopting conformations observed in the 8 nt mode. In human RPA, this linker is shorter (13 amino acids) compared to *U. maydis* RPA (28 amino acids). However, the residues in the first β-turn are conserved (G435,V436,G437,G438) (41). Our data likely reflects this structural transition from a flexible loop into a two β-turn upon binding to ssDNA. Furthermore, the loss in deuterium uptake amplitude upon DNA binding suggests occlusion of this region from solvent.

Upon DNA binding, major changes in the DBD-C structure are also observed around the Zn^2+^-finger motif. Four cysteines (C481, C486, C500, and C503) that coordinate zinc binding match peptides 30-32 and 34-35 (Figure 4f-h). Correspondingly, in our HDX analysis, peptides 32 to 37 have major changes in deuterium uptake (Figure 4f-j and Supplemental Figure 4b). The inner surface of DBD-C containing the DNA binding cleft shows significant decrease in deuterium exchange levels suggesting protection upon DNA binding and displays rapid exchange kinetics (peptides 32, 34; Figures 4f-g, and peptide 33; Supplemental Figure 4b). The decrease in D_2_O uptake reflects occlusion of this region from solvent upon DNA binding and the rapid kinetics denote direct DNA induced changes. Similarly, the solvent exposed outer surface region of the Zn^2+^-finger region showed a robust increase in deuterium uptake upon DNA binding with fast exchange kinetics (peptides 35, 36; Figures 4h-i).

Significant HDX is also observed in the helix-turn-helix motif (Supplemental Figure 5, HTH; α2a-α2b-α2c; 533 to 566 aa (4)), a region underneath the DNA binding cleft of DBD-C (peptides 38 to 43, Figures 4k-l, Supplemental Figures 4c-e). Peptide 37 is protected in presence of DNA, becoming more dynamic (exchange on a long timescale), having slow movements when DNA is absent (Figure 4j; **p=0.0011 at 3h). Upon DNA binding, the N-terminal half of HTH motif (peptides 38-40) showed reduction in deuterium uptake levels. The first incubation time point that shows statistical difference in deuterium uptake from the RPA alone are indicated for peptides 38-40 (p=ns, *p=0.0144, **p=0.0013) respectively (Figure 4k, and Supplemental Figures 4c & d). In contrast, the C-terminal half of HTH motif (peptides 41 to 43) displayed increased deuterium uptake and faster exchange kinetics (Figures 4l-m and Supplemental Figure 4e). The first time point post DNA addition that shows statistical difference in deuterium uptake are indicated for peptides 41-43 respectively (*p=0.0218, **p=0.0054, *p=0.0211) (Figures 4l-m, and Supplemental Figure 4e). Simultaneous protection and exposure in D_2_O exchange within this region suggests correlated conformational movements. That is, occlusion from solvent on the N-terminal side and exposure to solvent on the C-terminal side of HTH motif. It can be envisioned that DNA binding occludes the N-terminal end from deuteration by compaction which leads to conformational change that extends the C-terminal half. Indeed, in our modeled structure, we do see an extended conformation of HTH C-terminal half which can now contact DBD-D and DNA through *van der Waals* interactions (41)(Figure S5b). A similar interaction between DBD-C and DBD-D is observed in the cryoEM structure of S. cerevisiae RPA (23).

The L45 loop, connecting β-strand 4 and β-strand 5, is a flexible region in DBD-C and harbors Y581, a conserved residue that base-stacks with DNA. Another conserved aromatic residue F532 resides in β3-strand of DBD-C. Other residues in this region (R575, K577, Y581, and R586) also make extensive contacts with ssDNA (Supplemental Figure 5c) (42). Electron density for L45 in the structure of *U. maydis* Tri-C is poorly defined and suggests a dynamic region that is likely ordered upon binding to DNA (41). Peptide 44 corresponds to the L45 loop and is protected on DNA binding at early time points with a steady increase in HDX (Supplemental Figure 4f). Our data for DBD-C showcase robust and stable changes in deuteration levels upon interaction with DNA. In comparison to the highly dynamic FAB-half of RPA, DBD-C appears to stably interact with DNA with fewer dynamic changes.

We observe an interesting correlation in HDX patterns between peptide 37 in DBD-C and peptide 6 in PID^70N^. For both these peptides, DNA binding causes a change in slow HDX from 3 min onward post binding to DNA (compare Figure 3d and Supplemental Figure 4j). DNA binding causes deuterium uptake in peptide 6 compared to protection in DBD-C. Recent crosslinking experiments performed in our laboratories show physical interaction between these regions in the absence of DNA (Bothner and Antony, unpublished data). RPA can adopt many conformations given the flexible nature of the linkers. Thus, we propose a direct interaction between these two regions as one of the states and the interaction is perturbed upon DNA binding.

RPA32 harbors a disordered N-terminal region which is heavily phosphorylated, DBD-D, and PID^32C^. 17 total peptides for RPA32 were identified and analyzed: Peptides 1 to 5 cover the N-terminal region and show small changes in HDX upon DNA binding. Peptides 1 and 5 show rapid DNA binding dependent changes whereas peptides 2 and 4 do not. These data suggest that this region is undergoing conformational changes upon DNA binding, but it is interesting to note that this region is not part of the DNA binding domain in the structures. Peptides 9 to 13 encompass DBD-D and correspondingly show fast exchange kinetics at early time points, indicative of direct DNA binding induced changes in the structure. The data are consistent with DNA dependent occlusion/compaction of the DNA binding site (Figure 5 and Figure S6). One key structural motif in DBD-D is the β4-β5 hairpin (i.e. β4-L45-β5) which is an extension of the DNA binding site and displays rapid exchange (peptide 9 and 10) with a two-phase deuterium exchange profile observed for the β5 peptide (peptide 10). Upon DNA binding the C-terminal α-helix shows exposed (peptide 11) and protected (peptide 12) profiles (Figures 5h-i). These changes suggest that the α-helical bundle of Tri-C is dynamic and propagates DNA-induced conformational changes.

Interestingly, PID^32C^ (peptide 15, 16, 17) and its linker region (peptide 13) also show changes in deuteration in the presence of DNA (Figures 5j-l and Supplemental Figure 6c). Peptide 13 corresponds to the linker connecting DBD-D and PID^32C^ (Figures 2b region vi and 5a) and shows a significant (*p=0.0138) increase in deuterium uptake is observed suggesting solvent exposure (Figure 5j). One possibility is that interactions exist between PID^32C^ and DBD-D are perturbed upon DNA binding, a scenario similar to denoted above between PID^70N^ and DBD-C. Altered uptake patterns are observed in the tail-end of the helix in DBD-D (described above) and the linker. The deuteration profile of the PID^32C^ peptides do not show robust changes (peptide 15-17, Figures 5k-l, and Supplemental Figure 6c) similar to the observations for PID^70N^ (Figures 3b-e). An overlay of apo (PDB:1L10) and DNA bound (PDB:4GNX) DBD-D structures reveals mild changes (Supplemental Figure 6d). Thus, we conclude that both the protein interaction domains in human RPA do not interact directly with ssDNA.

**Figure 6.**
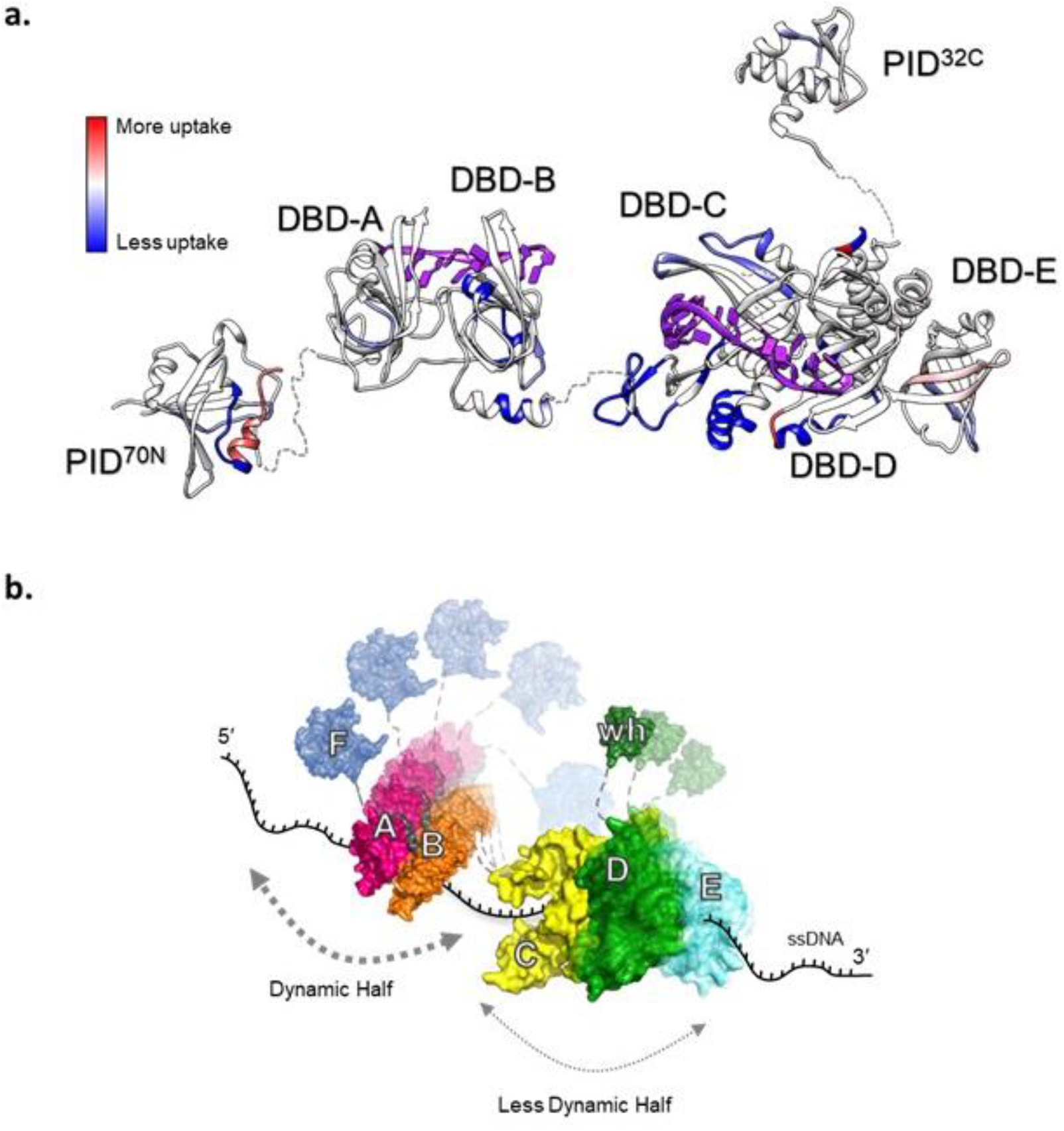
RPA binds to ssDNA using dynamic and less dynamic halves. **a**. HDX changes in human RPA upon binding to DNA are mapped onto a composite structure. Blue and red denote the scale of difference in deuterium uptake in the unbound versus DNA-bound forms of RPA. DNA binding primarily drives protection of regions from deuterium uptake as they become occluded from solvent. b. A dynamics-based model for RPA-DNA interactions is depicted. RPA is proposed to function as two halves. PID^70N^ (F domain), DBD-A and DBD-A form the dynamic half and have short residence on time. DBD-C, RPA32 (DBD-D) and RPA14 (DBD-E) form the less dynamic half and have longer residence time on DNA owing to the stability of DNA binding arising from the larger DBD-C domain. This model allows RPA to be stably bound to DNA while providing access to both ends of the DNA due to the dynamics of the individual DBDs. PID^70N^ and PID^32C^ primarily serve to bind and recruit RPA-interacting proteins onto DNA.

DBD-E is the third component of Tri-C. In the crystal structures, no direct interactions are observed between DNA and DBD-E. However, crosslinking experiments have suggested DNA interactions in this domain (6). Interestingly, in the cryoEM structure of multiple yeast RPA molecules on ssDNA, DBD-A of one RPA molecule interacts with DBD-E of the neighboring RPA. This interaction is further stabilized upon phosphorylation at position S178 in the FA-linker closer to DBD-A (23). Thus, DBD-E is an integral component of cooperative binding of RPA molecules. Eleven peptides were tracked in DBD-E (Supplemental Table 1). Our data show both *fast* (peptides 5-6, 8, 10) and *slow* (peptides 3-4, 7, 9, 10-11) D_2_O exchange for DBD-E. Several of the corresponding peptides (4-6, and 8-9) show DNA dependent solvent exposure (Figures 5m-q and Supplemental Figure 7). Peptides 3 and 4 are in the β1-strand region, significant (*p=0.017) exposure was observed for peptide 4 (Figure 5m). Peptide 5 is part of L34 loop in DBD-E and displays significant (*p=0.04) exposure kinetics at 3h (Figure 5n). There are two overlapping peptides for the β4 −β5 hairpin; where peptide 6 shows rapid and robust deuterium uptake (**p=0.001) while significant change was observed for peptide 7 (Figure 5o and Supplemental Figure S7b). This suggests solvent exposure of the β4 and L45 regions in DBD-E. Analogous regions in other DBDs (DBD-B peptides 18-19; DBD-C peptides 41-44; DBD-D peptides 9-10) show large conformational changes upon DNA binding. The loop between β5 and α2 region in DBD-E was covered by peptides 8, 9 and 10. Peptide 8 that overlaps both 9 and 10 showed significant (*p=0.03) and delayed deuterium uptake (Figure 5p). Interestingly, peptides 9 (*p=0.018) and 10 (p=ns) showed greater exchange reflecting exposure and protection respectively (Figure 5q and Supplemental Figure 7c). Thus, the entire DBD-E is exposed and the beginning of α2 (peptide 10) is protected (Figure 2f and Supplemental Figure 7c). However, most of the α2-helical region in DBD-E (peptide 11) is dynamic without any significant change in deuteration on DNA binding (Supplemental Figure 7d). The dynamic characteristics of the C-terminal α-helix in DBD-E is similar to the C-terminal α-helix in DBD-D (Figures 5h-i). Both these α-helices are part of the α-helix bundle in Tri-C.

Taken together, our data show that the Tri-C region of RPA is stable compared to the FAB region. Tri-C region of RPA shows rapid deuterium uptake in the presence of DNA and the deuteration levels off suggesting that the structures (H-bonding networks) are stable compared to the FAB region. Much of this stability arises from the interaction of DBD-C and DNA, with dynamic changes observed in DBD-D and DBD-E. Thus, within Tri-C, DBD-D and DBD-E are relatively dynamic than DBD-C but are less dynamic compared to DBD-A and -B. Our data support a model for RPA where DBDs-A and -B form the ‘*dynamic half*’ and DBDs-C, -D and -E form the ‘*less dynamic half*’ of RPA.

## DISCUSSION

One of the primary roles of RPA is to protect ssDNA from nucleases and form a nucleoprotein filament that serves to recruit other proteins. This is an inherently dynamic process in which RPA hands off the DNA to the incoming enzyme. The flexibility in RPA structure arises from the compartmentalization of its DBDs and PIDs which are tethered together by flexible linkers. The modular architecture allows the domains of RPA to employ both independent and coordinated DNA interactions that are tailored for specific functional outcomes. The formation of such conformations are further regulated by post-translational modifications (43).

Canonical models for RPA-DNA interactions primarily arose from biochemical investigations of isolated or select pairs of DBDs. DBDs-A and -B are defined as high affinity DNA binding domains. Tri-C is considered the weaker DNA binding portion of RPA (25). Recent single molecule and structural data suggest a dynamics-based picture of DBD-DNA interactions. DBD-A and DBD-B dissociate from DNA and interact with DBD-E from the neighboring molecule when multiple RPA molecules are visualized using cryoEM (23). The Tri-C is more ordered in these structures as well. In single molecule experiments where the DNA binding dynamics of individual DBDs are monitored, DBD-A and DBD-D can form four distinct conformations on DNA and dynamically transition between these states (25). In the presence of an RPA interacting protein, one of these states are not formed. Thus, a model was proposed wherein the dynamics of individual DBDs provides a mechanism to access RPA-bound ssDNA whilst not completely dissociating the RPA molecules. The single molecule experiments do not provide such information for the other DBDs or PIDs (due to technical limitations). Thus, a complete picture of the dynamics of all the domains of RPA and how they are rearranged in the presence of DNA is lacking.

The HDX data presented here provide a coarse-grain structural and kinetic landscape of the DNA-driven conformational changes (Figure 6a). The HDX profiles of all five DBDs (A, B, C, D, E) have altered exchange kinetics on a fast timescale in the presence of DNA. A result consistent with rapid on-rates for RPA-DNA binding. However, the exchange patterns of the HDX signal varies for the individual DBDs. The HDX of DBDs-A and -B rapidly re-equilibrate suggesting rapid on-off kinetics. Thus, these domains and their interactions are dynamic in nature as denoted by the early timepoints in the HDX analysis. DBDs-C, -D and -E also show large scale deuterium uptake profiles and show slow re-equilibration. HDX data at the later timepoints reveal that deuterium uptake levels off over a period of 24 hours for many of their peptides suggesting formation of stable H-bonding networks (Figures 4 and 5). Thus, the DBDs in the Tri-C region are less dynamic compared to the ones in the FAB half of RPA.

We do not observe fast exchange kinetics or robust deuterium changes in either PID^70N^ or PID^32C^ suggesting that the primary roles of these domains are to interact with RIPs. Binding of p53 to the cleft in PID^70N^ has been shown to be outcompeted by DNA, and thus proposed to possess very weak DNA binding properties. But more importantly, PID^70N^ serves as a high affinity binding site for more than two dozen proteins including p53 and Dna2 (44)(45). In addition, PID^70N^ interacts with the N-terminal phosphorylation motif of RPA32 (46), and preliminary crosslinking-MS data suggest an interaction between PID^70N^ and DBD-C (Bothner and Antony, unpublished work). Thus, the HDX changes we observe in the later timescales likely originate from the release of such intra-RPA:PID^70N^ interactions upon DNA binding to DBD-C and DBD-D, and not from direct DNA binding to PID^70N^.

Biophysical characterization of full-length RPA suggests that DNA binding is highly dynamic and that the interaction of individual DBDs is not sequential as previously envisioned (23)(25)(47)(48)(49). The correlated movements/dynamics of the DBDs (and likely PIDs) enable rapid RPA diffusion on ssDNA (∼5,000 nt^2^ sec^−1^ at 37 °C) and efficiently melt small hairpins and duplex regions as it diffuses (7). RPA–DNA complexes have also been shown to exchange rapidly in the presence of free RPA (26)(48). Transition between specific DNA binding modes are thought to promote such mechanical movements of RPA. An initial 8-10 nt is formed first followed by transition to a 20 nt mode and finally resulting in the 30 nt mode where all DBDs are DNA-bound (1)(13)(50). RPA dynamically forms at least two kinetically distinct complexes on DNA with different stabilities that are microscopically dissociating and re-associating (47).

Our data support a DNA binding dynamics-based model for RPA-DNA interactions. We envision RPA to function as two dynamic halves. The FAB half and the Tri-C half (Figure 6b). The FAB half is *dynamic* with rapid on-off kinetics on ssDNA and the Tri-C half is *less-dynamic* with more stable interactions with DNA, primarily driven by the stability of DBD-C. The HDX data do not support a role for the PIDs in mediating direct DNA interactions, but intra-domain interactions exist between the PIDs and the DBDs. These interactions are released as conformational changes are driven upon DNA binding as slow exchange HDX is observed for both PIDs. The dynamics-based model presented here would allow RPA to maintain high affinity interactions through stable binding by DBD-C and contributions from other parts of Tri-C. The dynamic interactions of DBDs-A and -B would enable RIPs that interact with PID^70N^ to gain access to the 5′ end of the RPA coated DNA. Similarly, even the less-dynamic DBDs-D and -E can be remodeled by RIPs that interact with PID^32C^ and bind to the 3′ of the RPA nucleoprotein filament as previously observed for Rad52 (25). When multiple RPA molecules are bound on the DNA, the dynamic nature of the DBDs would enable RIPs to bind to the DNA.

## SUPPLEMENTARY DATA

Supplementary Data are available at NAR online.

## FUNDING

This work was supported by grants from the National Institute of General Medical Sciences of the National Institutes of Health to E.A. (R01GM130746 and R01GM133967). Funding for the Mass Spectrometry facility used in this publication was made possible in part by the MJ Murdock Charitable Trust and the National Institute of General Medical Sciences of the National Institutes of Health (P20GM103474). Funding for open access charge: National Institutes of Health.

## CONFLICT OF INTEREST

The authors declare no conflict of interests.

## Notes

### Competing Interest Statement

The authors have declared no competing interest.

