## Supplemental Information for "Hydrogen-deuterium exchange reveals a dynamic DNA binding map of Replication Protein A"

Pepsin generated RPA peptides identified in HDX-MS

**RPA70 ± d(T)$_{30}$ oligonucleotide**

| Peptide[a] | Residues[b] | Sequence |
|---|---|---|
| 1 | 1-25 | MVGQLSEGAIAAIMQKGDTNIKPIL |
| 2 | 12-25 | AIMQKGDTNIKPIL |
| 3 | 13-25 | IMQKGDTNIKPIL |
| 4 | 45-56 | LMSDGLNTLSSF |
| 5 | 46-58 | MSDGLNTLSSFML |
| 6 | 67-73 | EEEQLSS |
| 7 | 88-92 | KDGRR |
| 8 | 246-256 | FPLIEVNKVYY |
| 9 | 247-257 | PLIEVNKVYYF |
| 10 | 298-301 | VQFD |
| 11 | 297-307 | TVQFDFTGIDD |
| 12 | 301-308 | DFTGIDDL |
| 13 | 301-316 | DFTGIDDLENKSKDSL |
| 14 | 305-316 | IDDLENKSKDSL |
| 15 | 349-357 | MDTSGKVVT |
| 16 | 361-376 | WGEDADKFDGSRQPVL |
| 17 | 365-376 | ADKFDGSRQPVL |
| 18 | 395-411 | SSSTIIANPDIPEAYKL |
| 19 | 395-415 | SSSTIIANPDIPEAYKLRGWF |
| 20 | 409-415 | YKLRGWF |
| 21 | 416-426 | DAEGQALDGVS |
| 22 | 416-429 | DAEGQALDGVSISD |
| 23 | 416-434 | DAEGQALDGVSISDLKSGG |
| 24 | 427-445 | ISDLKSGGVGGSNTNWKTL |
| 25 | 431-444 | KSGGVGGSNTNWKT |
| 26 | 431-446 | KSGGVGGSNTNWKTLY |
| 27 | 446-461 | YEVKSENLGQGDKPDY |
| 28 | 447-464 | EVKSENLGQGDKPDYFSS |
| 29 | 455-463[c] | QGDKPDYFS |
| 30 | 477-485[c] | MYQACPTQD |
| 31 | 479-497[c] | QACPTQDCNKKVIDQQNGL |
| 32 | 486-497 | CNKKVIDQQNGL |
| 33 | 487-497 | NKKVIDQQNGL |
| 34 | 498-510 | YRCEKCDTEFPNF |
| 35 | 502-504 | KCD |
| 36 | 502-511 | KCDTEFPNFK |
| 37 | 511-516 | KYRMIL |
| 38 | 536-542 | AEAILGQ |
| 39 | 541-558 | GQNAAYLGELKDKNEQAF |
| 40 | 552-565 | DKNEQAFEEVFQNA |
| 41 | 558-667 | FEEVFQNANF |
| 42 | 559-564 | EEVFQN |
| 43 | 559-567 | EEVFQNANF |
| 44 | 579-585 | ETYNDES |
| 45 | 591-606[c] | VMDVKPVDYREYGRRL |

**RPA32 ± d(T)$_{30}$ oligonucleotide**

| Peptide[a] | Residues[b] | Sequence |
|---|---|---|
| 1 | 1-6 | MWNSGF |
| 2 | 5-16 | GFESYGSSSYGG |
| 3 | 15-22[c] | GGAGGYTQ |
| 4 | 15-28 | GGAGGYTQSPGGFG |
| 5 | 23-39 | SPGGFGSPAPSQAEKKS |
| 6 | 63-66[c] | VFRI |
| 7 | 107-111 | WVDTD |
| 8 | 107-124 | WVDTDDTSSENTVVPPET |
| 9 | 126-129 | VKVA |
| 10 | 142-144 | VAF |
| 11 | 166-169 | HMVL |
| 12 | 166-190 | HMVLSKANSQPSAGRAPISNPGMSE |
| 13 | 190-197 | EAGNFGGN |
| 14 | 193-214[c] | NFGGNSFMPANGLTVAQNQVLN |
| 15 | 216-227 | IKACPRPEGLNF |
| 16 | 221-225 | RPEGL |
| 17 | 267-276 | TDAEHHHHHH |

**RPA14 ± d(T)$_{30}$ oligonucleotide**

| Peptide[a] | Residues[b] | Sequence |
|---|---|---|
| 1 | 1-27[c] | MVDMMDLPRSRINAGMLAQFIDKPVCF |
| 2 | 11-14[c] | RINA |
| 3 | 28-40 | VGRLEKIHPTGKM |
| 4 | 29-40 | GRLEKIHPTGKM |
| 5 | 60-62 | DEE |
| 6 | 65-82 | GIVEVVGRVTAKATILCT |
| 7 | 72-81 | RVTAKATILC |
| 8 | 88-98 | KEDSHPFDLGL |
| 9 | 89-95 | EDSHPFD |
| 10 | 95-97 | DLG |
| 11 | 98-108 | LYNEAVKIIHD |

*a* Peptides are numbered sequentially
*b* Peptide sequence length
*c* Some data missing, indicated 'white box' in heatmap Fig. 2A-C, and uptake curve is not displayed for them. Additionally, peptide RPA70;WT *α*27(24h, 3h;n=2), and *α*28(0.05h;n=2) had indicated experimental replicated.

**Supplemental Table 1. Table 1. Pepsin generated RPA peptides identified in HDX experiments.** A list of peptides obtained digesting RPA with the protease pepsin and corresponding sequence. Peptides are numbered sequentially as '*a*'; Peptide sequence length indicated as '*b*'. Data point not available '*c*'. Residues are color coded as in Figure 1a. *a;* Peptides are numbered sequentially. *b; Peptide* sequence length. *c:* Some data missing, indicated in heatmap Figure 2a-2c, and uptake curve is not displayed for them. Additionally, peptide RPA70; apo RPA *a*27(24h,3h; n=2), and *a*28(0.05h; n=2) had indicated experimental replicates. Peptides sequence are color coded to match the crystal structure as in Figure 1a; purple (DBD-F/PID$^{70N}$), pink (DBD-A), brown (DBD-B), yellow (DBD-C), red (Pd motif), light green (DBD-D), dark green (wh/ PID$^{32C}$), cyan (DBD-E/RPA14), and grey (Linker).

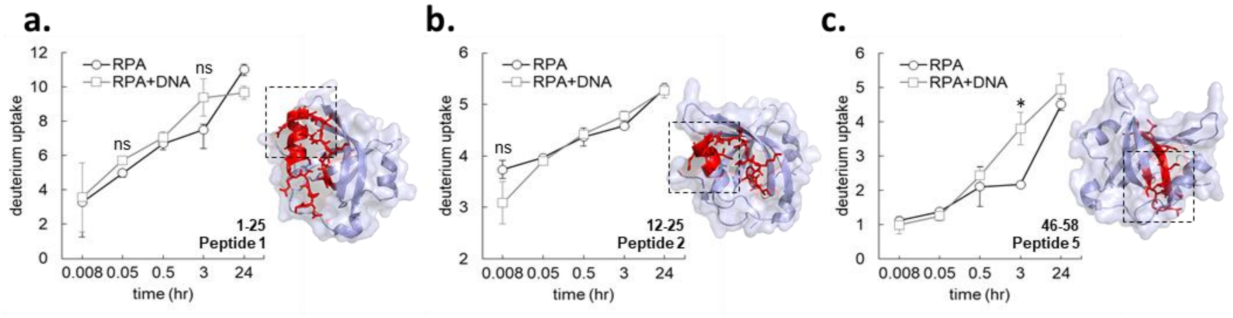


**Supplemental Figure 1**. **HDX-MS data corresponding to PID$^{70N}$ (F-domain).** Deuterium uptake data for peptides identified in the HDX-MS analysis for PID$^{70N}$ are shown. Each peptide is shown in red in the corresponding structure of PID$^{70N}$.

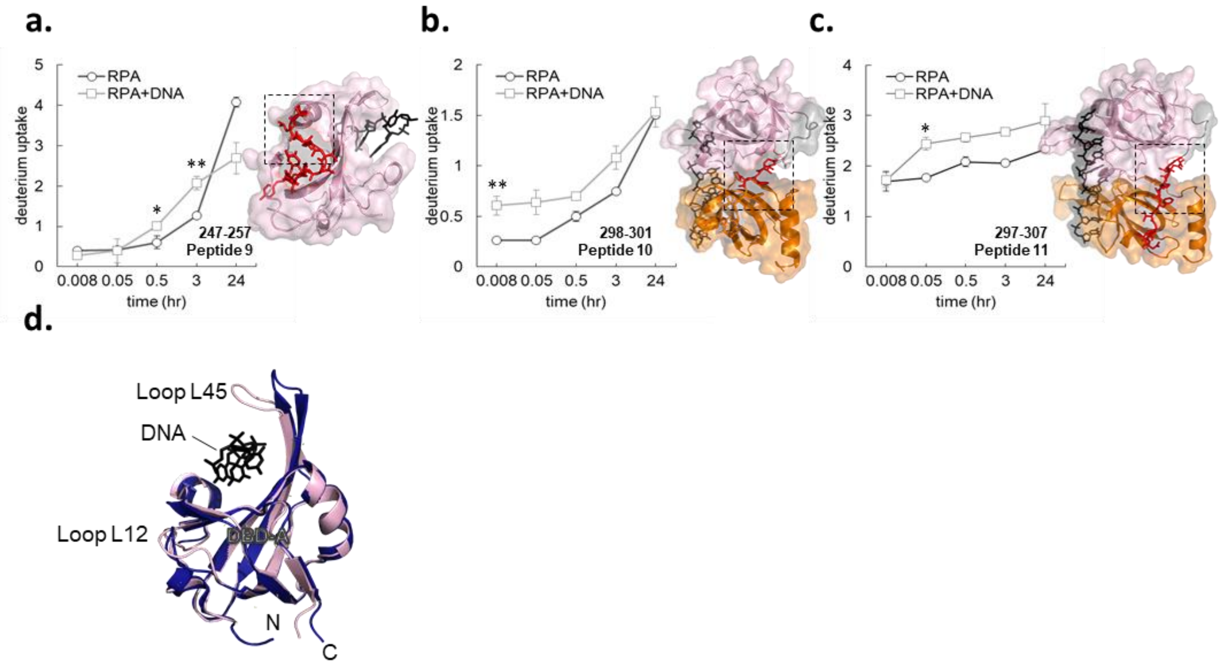


**Supplemental Figure 2**. **HDX-MS data corresponding to DBD-A and DBD-B in RPA 70. a-d.** Deuterium uptake data for peptides identified in the HDX-MS analysis for DBD-A and some peptides in DBD-B are shown. Each peptide is shown in red in the corresponding structure of DBD-A (fusia) and DBD-B (orange). DNA is shown as black sticks. **e.** An overlay of the X-ray crystal structures of the apo DBD-A from Homo Sapiens (PDB:1FGU; blue) and DNA-bound DBD-A from Homo sapiens (PDB:1JMC; pink) highlight the DNA-induced conformational changes. The largest changes are observed in loops L45 and L12.

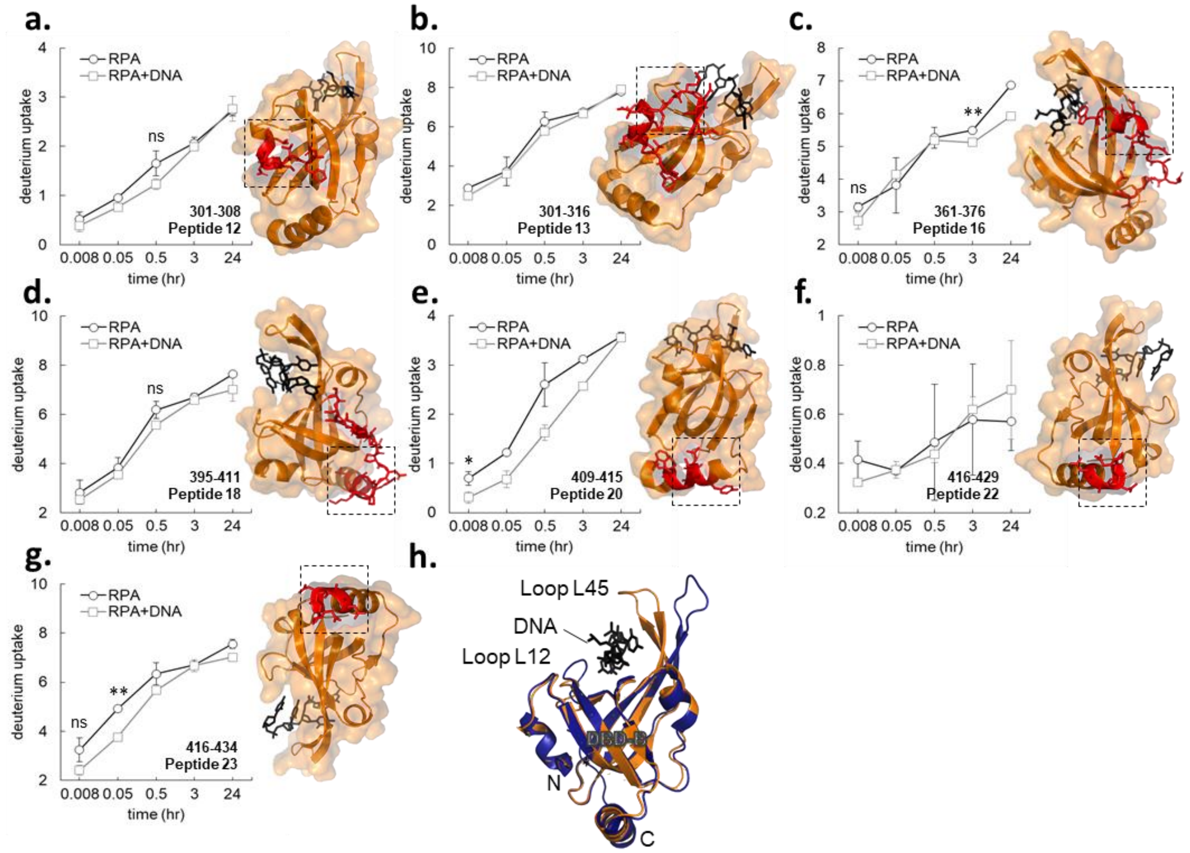


**Supplemental Figure 3**. **HDX-MS data corresponding to DBD-B in RPA 70. a-g.** Deuterium uptake data for peptides identified in the HDX-MS analysis for DBD-B are shown. Each peptide is shown in red in the corresponding structure of DBD-B (orange). DNA is shown as black sticks. **h.** An overlay of the X-ray crystal structures of the apo DBD-B from Homo sapiens (PDB:1FGU; blue) and DNA-bound DBD-B from Homo sapiens (PDB:1JMC; brown) highlight the DNA-induced conformational changes. The largest changes are observed in loops L45 and L12.

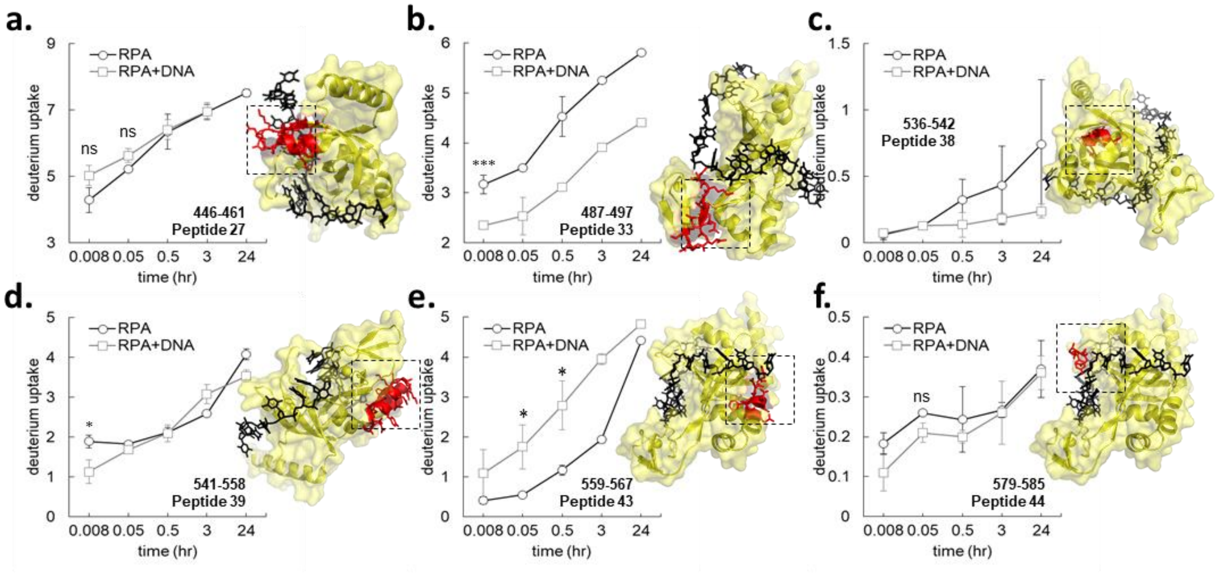


**Supplemental Figure 4**. **HDX-MS data corresponding to DBD-C in RPA 70. a-f.** Deuterium uptake data for peptides identified in the HDX-MS analysis for DBD-C are shown. Each peptide is shown in red in the corresponding structure of DBD-C (orange). DNA is shown as black sticks. Since a DNA-bound structure of DBD-C is not available, we used SWISS MODEL to generate the structure. The DNA-bound DBD-D structure from *Ustilago maydis* RPA (4GNX) was used as a template to generate the model.

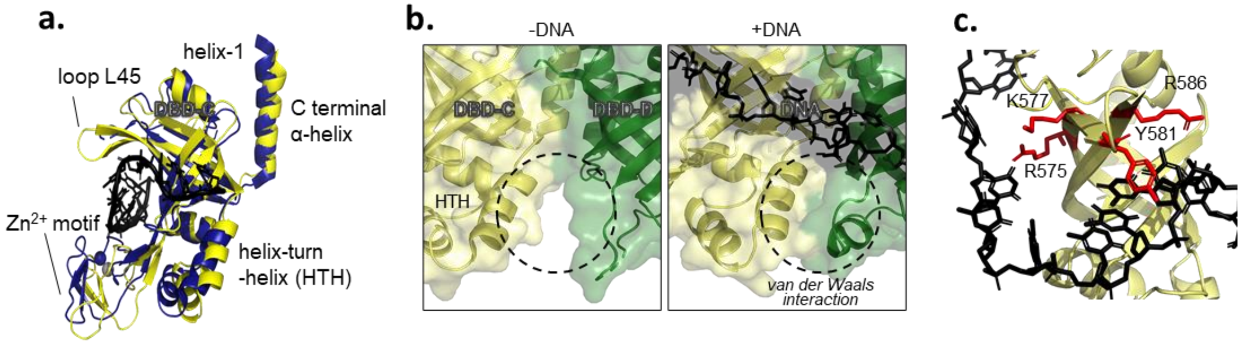


**Supplemental Figure 5. Comparison of apo and DNA-bound structures of DBD-C in RPA70. a.** Overlay of the X-ray crystal structures of the apo DBD-C from Homo sapiens (PDB:1L1O; blue) and DNA bound DBD-C (SWISS MODEL using *Ustilago maydis* PDB:4GNX; yellow) highlights the extensive conformational changes. Changes in the $Zn^{2+}$-finger domain, loop L45, the helix-turn-helix motif, and the C-terminal a-helix are observed upon binding to ssDNA. **b.** Van Der Waals contacts between DNA and the HTH motif are depicted. The change in compaction upon DNA binding is denoted by the dotted region. **c.** Residues in the L45 loop of DBD-C that make direct contacts with the ssDNA are shown as red sticks.

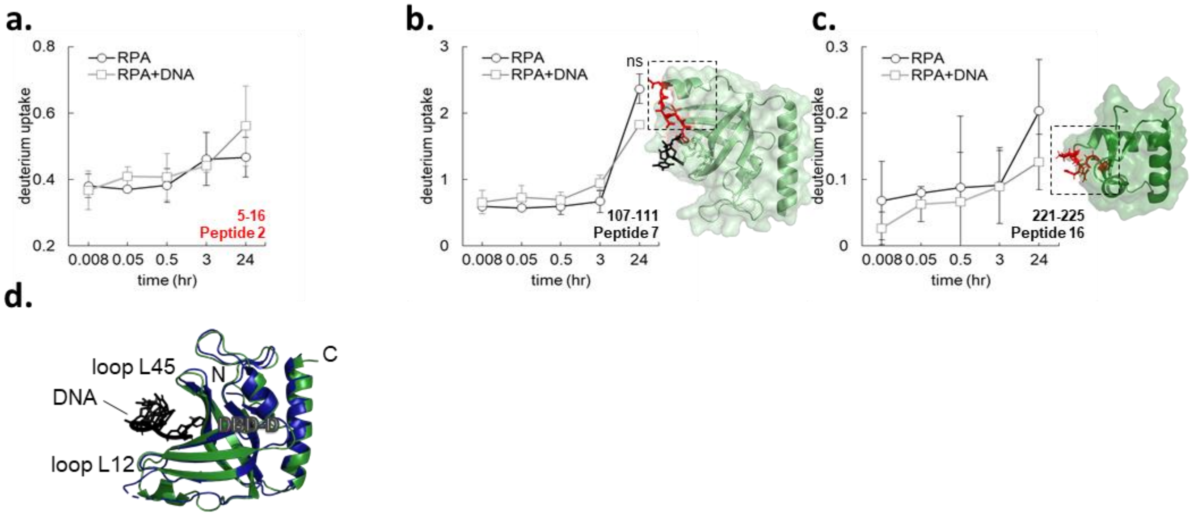


**Supplemental Figure 6**. **HDX-MS data corresponding to DBD-D in RPA32. a-c.** Deuterium uptake data for peptides identified in the HDX-MS analysis for DBD-D are shown. Each peptide is shown in red in the corresponding structure of DBD-D (green). DNA is shown as black sticks. **d.** Overlay of the X-ray crystal structure of the apo DBD-D from Homo sapiens (PDB:1L1O; blue) and DNA-bound DBD-D (SWISS-MODEL using *Ustilago maydis* PDB:4GNX; green) are shown. Conformational changes in Loops L12 and L45 are observed in the presence of DNA.

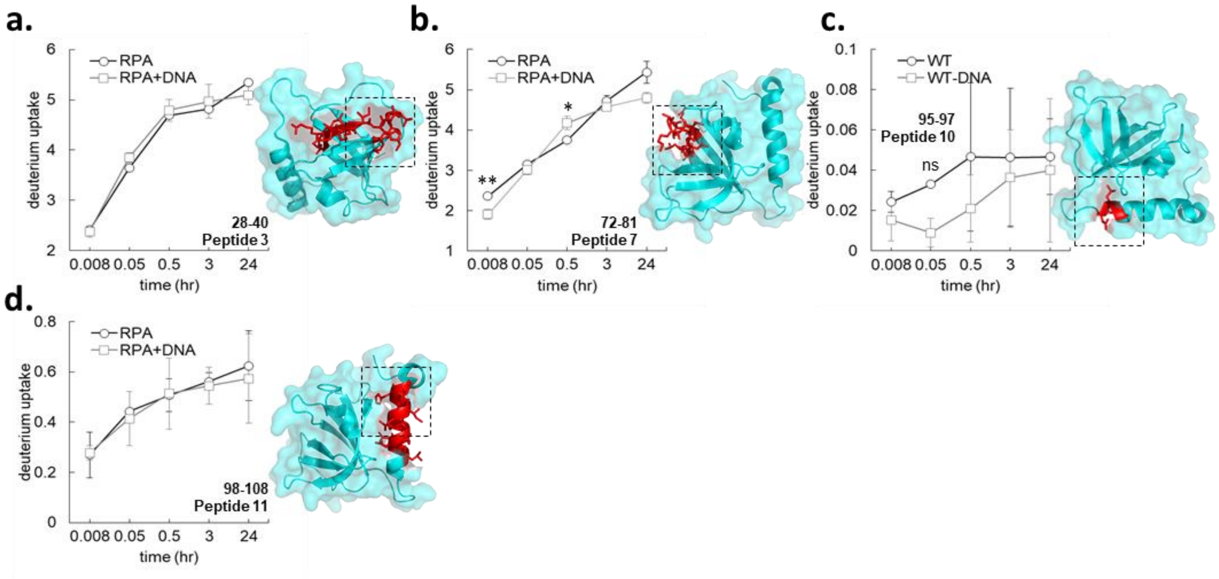


**Supplemental Figure 7**. **HDX-MS data corresponding to DBD-E in RPA14.** Deuterium uptake data for peptides identified in the HDX-MS analysis for DBD-E in RPA14 are shown. Each peptide is shown in red in the corresponding structure of RPA14 (cyan).
